# Single-Domain Antibodies as Potent Inhibitors of Clinically Relevant β-Lactamases in Multidrug-Resistant Bacteria

**DOI:** 10.64898/2025.12.03.691997

**Authors:** Paloma Osset-Trénor, Markus Proft, Amparo Pascual-Ahuir

**Affiliations:** Centro de Investigación e Innovación en Bioingeniería Ci2B, Universitat Politècnica de València, Ciudad Politécnica de la Innovación, Edificio 8B, Camino de Vera s/n, 46022 Valencia, Spain; Department of Metabolism, Inflammation and Aging, Instituto de Biomedicina de Valencia IBV-CSIC, 46010 Valencia, Spain; Valencia Biomedical Research Foundation, Centro de Investigación Príncipe Felipe (CIPF) – Associated Unit to the Instituto de Biomedicina de Valencia IBV-CSIC, Valencia, Spain

## Abstract

Antimicrobial resistance (AMR) represents a critical threat to global health, largely driven by the dissemination of β-lactamases that inactivate frontline antibiotics. Among the most problematic are BlaMab-2 from *Mycobacterium abscessus*, KPC-2 and OXA-48 from *Klebsiella pneumoniae*, and VIM-2 from *Pseudomonas aeruginosa*, which together confer broad resistance to β-lactams and carbapenems. Current β-lactamase inhibitors face declining efficacy as resistance variants continue to emerge, underscoring the need for innovative strategies.

Here, we explored single-domain antibodies (sd-Abs) as enzyme-directed inhibitors of β-lactamases. A library of sd-Abs was screened, and two candidates, B2 and B5, were characterized *in vitro* and *in vivo*. Both sd-Abs inhibited BlaMab-2 activity in *E. coli* expression systems, following a competitive inhibition mechanism, with B2 consistently displaying stronger potency (K_i_ ≈ 1.5 µM) than B5. Remarkably, B2 also demonstrated broad inhibitory activity against KPC-2, VIM-2, and OXA-48, while B5 showed an alternative inhibition profile, including uncompetitive characteristics against VIM-2 and OXA-48. Comparison with clinically deployed inhibitors revealed that the K_i_ values of B2 and B5 are of the same order of magnitude—or superior in some cases—highlighting their therapeutic promise.

Our findings establish sd-Abs as a versatile platform for the inhibition of diverse β-lactamases, with B2 emerging as the most broadly effective candidate. By expanding the utility of existing β-lactams, sd-Abs could help restore antibiotic efficacy against multidrug-resistant pathogens. This study underscores the potential of antibody-based enzyme inhibitors as a new class of anti-resistance therapeutics.

## Introduction

Antibiotic resistance is one of the most urgent global health challenges of our time. A landmark study published in *The Lancet* estimated that nearly 5 million deaths per year are associated with antibiotic-resistant bacterial infections, with the highest burden in low- and middle-income countries. This series also highlighted how this threat affects individuals across the entire life course, with newborns, older adults, and people with chronic conditions being particularly vulnerable (Murray et al., 2022). Beyond the health impact, antimicrobial resistance (AMR) threatens to impose a catastrophic economic toll: at current rates of inaction, treatment of resistant infections alone is projected to cost $412 billion per year by 2035, while losses in labour productivity may exceed $443 billion annually (Global Leaders Group on Antimicrobial Resistance, 2024).

β-lactam antibiotics are the most widely prescribed class of antibacterial agents due to their broad-spectrum activity. These compounds are characterised by a four-membered β-lactam ring, that covalently binds to penicillin-binding proteins (PBPs), thereby inhibiting the final steps of peptidoglycan cross-linking in the cell walls of Gram-negative and Gram-positive bacteria (Blumberg & Strominger, 1974; Tipper & Strominger, 1965). Although bacteria employ multiple mechanisms to evade antibiotic action, the production of β-lactamases remains the most prevalent and clinically significant, especially in Gram-negative pathogens. These enzymes catalyse the hydrolysis of the β-lactam ring of penicillins, cephalosporins, monobactams and carbapenems, nullifying their antibacterial efficacy.

According to Ambler’s classification, which is based on sequence homology and fundamental differences in the hydrolytic mechanism, β-lactamases are divided into four classes: A, B, C, and D. Specifically, classes A, C, and D share the presence of a serine residue at the enzyme’s active site (serine β-lactamases; SBL), whereas class B consists of a heterogeneous group of zinc-dependent metalloenzymes (metallo-β-lactamases or MBL). These four classes are widely distributed, however, certain enzyme families have become especially prevalent among clinically significant pathogens, mainly Gram-negative bacteria responsible for opportunistic infections associated with medical care of immunocompromised patients. These key enzyme families include TEM, SHV, CTX-M, and KPC (class A); NDM and VIM (class B); and CMY and ADC (class C). Class D enzymes are called oxacillinases (OXA) and include the OXA-23 and 24/40 groups, as well as OXA-48, which are particularly concerning due to their role in carbapenem resistance in *Acinetobacter baumannii* and *Enterobacteriaceae*, respectively (Tooke et al., 2019).

Our study focuses on representatives of the three most clinically relevant β-lactamase families: class A (BlaMab-2 from *Mycobacterium abscessus* and KPC-2 from *Klebsiella pneumoniae*), class D (OXA-48 from *Klebsiella pneumoniae*), and class B (VIM-2 from *Pseudomonas aeruginosa*) (Figure 1). *Mycobacterium abscessus* is a non-tuberculous mycobacterium (NTM) ranked as the second most important pathogen in NTM-associated pulmonary diseases (Cristancho-Rojas et al., 2024). Its broad resistance to antibiotics is largely attributed to its low-permeability cell envelope (Nguyen et al., 2024) and the production of the BlaMab β-lactamase (Dubée et al., 2014), whose deletion increases susceptibility to β-lactams such as amoxicillin and ceftaroline (Lefebvre et al., 2016). *Klebsiella pneumoniae* and *Pseudomonas aeruginosa* are Gram-negative pathogens listed by the World Health Organization as “critical priority” for new antibiotic development (World Health Organization, 2024). In *K. pneumoniae*, resistance is partly mediated by carbapenemases such as KPC-2 (*Klebsiella pneumoniae* carbapenemase) and the carbapenem-hydrolysing oxacillinase OXA-48 (Poirel et al., 2004). *P. aeruginosa* produces the Verona integron–encoded metallo-β-lactamase VIM-2, which efficiently hydrolyses carbapenems (Fortunato et al., 2023), and it is further distinguished by its capacity to form biofilms, significantly complicating treatment.

**Figure 1.**
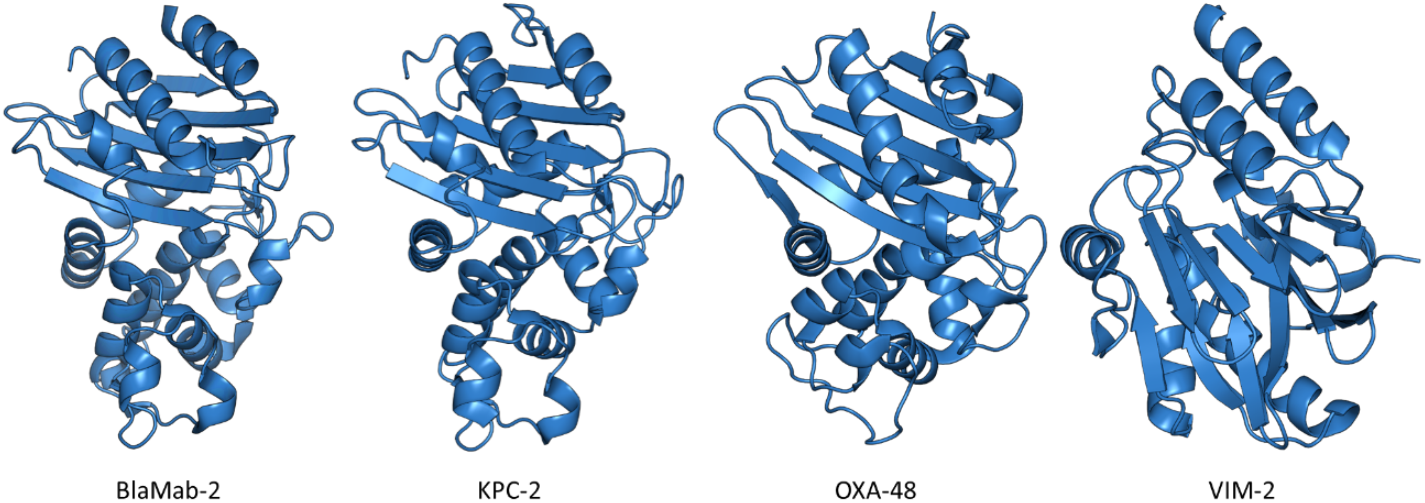
Structural representations of clinically relevant β-lactamases targeted in this study. Cartoon set specular, off models of the β-lactamase enzymes BlaMab-2 (*Mycobacterium abscessus*, PDB ID: 4YFM), KPC-2 (*Klebsiella pneumoniae*, PDB ID: 5UL8), OXA-48 (*Klebsiella pneumoniae*, PDB ID: 5DTK), and VIM-2 (*Pseudomonas aeruginosa*, PDB ID: 4BZ3) selected as representatives of Ambler classes A, D and B. Structures were visualized in PyMOL using a uniform color scheme and white background. Only one chain is shown per enzyme for clarity.

Collectively, these organisms represent important nosocomial pathogens, capable of causing opportunistic infections in immunocompromised patients, and exhibit multidrug resistance profiles. Their capacity to evade multiple lines of antimicrobial therapy represents a growing challenge in intensive care units and in patients with significant comorbidities.

Given the escalating threat of AMR, it is crucial to develop novel therapeutic strategies that either directly combat resistant bacteria or enhance the efficacy of existing antibiotics. Phage therapy, which employs highly specific lytic bacteriophages to lyse pathogens without damaging the commensal microbiota (Viertel et al., 2014), has demonstrated efficacy in both preclinical models and clinical case series of refractory infections and biofilms (Gordillo Altamirano & Barr, 2019). Although its combination with antibiotics can produce synergistic effects and reduce the selection of resistant mutants (Comeau et al., 2007; Knezevic et al., 2013), phage therapy is also affected by bacterial resistance (Gordillo Altamirano & Barr, 2019). In parallel, phage-derived endolysins are hydrolases produced by phages that degrade the peptidoglycan wall of Gram-positive and Gram-negative bacteria with high specificity. In addition, they can be designed or fused with penetration peptides to act more efficiently (Schmelcher et al., 2012) and have established themselves as therapeutic candidates with potent bactericidal activities and a low resistance profile (Rahman et al., 2021). In this context, single-domain antibodies (sd-Abs) stand out for their small size (~15 kDa), high stability, and ability to be designed with high affinity and specificity towards critical enzyme targets. In addition, they can be obtained through directed evolution by random mutagenesis and selection from combinatorial libraries (through phage display or yeast surface display). These properties, combined with their efficient production in bacterial systems, facilitate their design as bacterial enzyme inhibitors, including the specific inhibition of β-lactamases. Thus sd-Abs are positioned as particularly promising candidates for combating multidrug-resistant bacteria, including those addressed in this study (Boder & Wittrup, 1997; Muyldermans, 2013).

Several β-lactamase inhibitors are co-administered with β-lactam antibiotics to enhance their efficacy. However, bacteria continuously evolve mechanisms to evade these inhibitors, leading to an increasing number of resistant strains. The aim of this study is to demonstrate the ability of two sd-Abs to neutralize β-lactamases from multidrug-resistant bacteria, specifically BlaMab-2, evaluate their potential as enzymatic inhibitors and determine their efficacy against other clinically relevant β-lactamases, including KPC-2, VIM-2 and OXA-48 from different bacterial species.

## Material and Methods

### Cloning

To obtain *E. coli* strains producing the four β-lactamases, synthetic DNA sequences for BlaMab-2 (PDB ID: 4YFM), KPC-2 (PDB ID: 5UL8), VIM-2 (PDB ID: 4BZ3), and OXA-48 (PDB ID: 5DTK) were obtained from Twist Bioscience. These sequences were amplified by PCR using the primers A (forward, 5’-CAATCCGCCCTCACTACAACCG) and and B (reverse, 5’-TCCCTCATCGACGCCAGAGTAG). The amplified products were cloned into the pET-26b(+) vector via enzymatic digestion using the *Bam*HI restriction enzyme and subsequently transformed into *E. coli* BL21(DE3) cells (*Thermo Scientific™ BL21(DE3) Competent Cells*), following the manufacturer’s protocol.

A collection of 96 sd-Abs was screened, and the seven best candidates (B1-B7) were selected based on their affinity and specificity. The DNA encoding for the sd-Abs was amplified by PCR using the primers C (forward, 5’-CCCTC**GGATCC**GCAGGTGCAGCTGCAGGAAAGCGGC) and D (reverse, 5’-CCCC**GGATCC**GAGCTGCTCACGGTCACCTGGGTGCC) which added *Bam*HI (bold) restriction sites. The cloning into the pET-26b(+) plasmid was confirmed by colony PCR and sequencing (performed by Macrogen) in kanamycin-resistant colonies.

### Expression and purification of recombinant proteins

For the production of BlaMab-2 and the sd-Abs, each recombinant strain (*E. coli* BL21(DE3) harbouring the respective plasmid) was grown in LB medium supplemented with 50µg/ml kanamycin at 37º overnight with agitation. The following day, the medium was changed to fresh LB medium and expression was induced with 1mM β-D-1-thiogalactopyranoside (IPTG) at 16°C overnight with agitation. For protein extraction, cells were centrifuged at 3400xg for 20 min and the pellet was resuspended in sucrose solution (10mM Tris-HCl pH 8, 1M EDTA, 25% (w/v) Sucrose) at five times the cell pellet weight. After 10 minutes incubation at room temperature, cells were centrifuged at 14000xg for 45 minutes. The resulting pellet was resuspended in a cold shock solution (10mM Tris-HCl pH 8, 0.5M MgCl_2_), incubated on ice for 10 minutes and centrifuged again at 14000xg for 25 minutes. The supernatant containing the protein fraction, was collected and stored at 4°C. The extraction step was repeated to maximize protein yield. Proteins were filtered through a 22μm pore filter and purified using affinity chromatography with a HisTrap^TM^ HP column (Cytiva). Proteins were eluted with 500mM imidazole. BlaMab-2 was stored in 50% glycerol at −20°C, while sd-Abs were stored at 4°C.

### Verification of protein production

The expression of the four β-lactamases was confirmed by sodium dodecyl sulphate polyacrylamide gel electrophoresis (SDS-PAGE) and Western blot analysis. For SDS-PAGE, 15% gels were used, and proteins were denatured by heating at 96°C in loading buffer (7.5% SDS, anti rb antibody 0.1M dithioerythritol, 10mM EDTA, 30% sucrose (w/v), 0.25mg/ml bromophenol blue and 0.3M Tris, adjusted to pH 6.8 with HCl). The PageRuler^TM^ Prestained Protein Ladder (Thermo Scientific) was used as a molecular weight marker. Electrophoresis was performed at 120 V in running buffer (192 mM glycine and 0.1 % SDS brought to pH 8.3 with Tris), followed by staining with BlueSafe protein staining reagent (NZYtech) for 15 min and destained with distilled water under gentle agitation.

For Western-Blot analysis, proteins were transferred from the SDS-PAGE gels onto PVDF membranes using the Bio-Rad Western-Blot transfer system at 30 V overnight. Membranes were blocked with 2% skimmed milk powder in Tris-buffered saline (TBS-M) for 30 minutes, then incubated with anti-His rabbit primary antibody (Jackson InmunoResearch Laboratories, Inc.) (1:1000 dilution in TBS-M) for 1 hour. After washing, membranes were incubated with anti-rabbit-ECL secondary antibody (Cityiva) (1:5000 dilution in TBS-L) for 30 minutes. Protein detection was performed using the ImageQuant 800 system.

### In vitro enzyme activity assays of BlaMab-2 and inhibition by B2, B5 and B7

The activity of purified BlaMab-2 was assessed using the chromogenic β-lactam substrate nitrocefin, a yellow chromogenic substrate that is hydrolysed by β-lactamases to produce a red product detectable at a wavelength of 490nm (Spicer et al., 2010). BlaMab-2 was incubated with increasing concentrations of nitrocefin (20 µM, 40 µM, 60 µM, 80 µM, 100 µM, and 200 µM) diluted in sterile milli-Q water. The protein was used at a concentration of 9mM and diluted 10^4^ times in phosphate buffer saline (PBS) plus 0.2% of BSA (PBSA) at pH 5.8. The hydrolysis reaction by BlaMab-2 was monitored by measuring absorbance at 490 nm every 60s using a TECAN Infinite 200 PRO microplate reader. For inhibition assays, BlaMab-2 (same concentration as in the previous assay) was incubated with the sd-Abs (B2, B5 and B7) at a final concentration of 2μM for 1 hour at 4°C under gentle agitation. Nitrocefin hydrolysis was then assessed as described above. Each assay was performed in triplicate.

### In vivo enzyme activity assays of BlaMab-2, KPC-2, VIM-2 and OXA-48 and their inhibition by B2 and B5

The enzymatic activity of BlaMab-2, KPC-2, VIM-2, and OXA-48 was evaluated in a cellular context using recombinant *E. coli* BL21(DE3) strains. Cells were grown to an OD_600_ of 0.6-0.8 and collected by centrifugation. The pellet was resuspended in PBSA at pH 7.4 and diluted 100-fold. Nitrocefin was added at increasing concentrations (20 µM, 40 µM, 60 µM, 80 µM, 100 µM, and 200 µM) diluted in sterile H_2_O milli-Q and the absorbance at 490 nm was measured every 45 seconds using a TECAN Infinite 200 PRO microplate reader. For *in vivo* inhibition assays, bacterial cells were incubated with B2 and B5 at a final concentration 20 µM for 1 hour at room temperature before adding nitrocefin. Enzyme activity was assessed as described above.

### Determination of kinetic parameters

The concentration of hydrolyzed nitrocefin (c) was calculated from absorbance (A), the molar extinction coefficient of hydrolyzed nitrocefin (ε = 15,900 M^−1^·cm^−1^, reported at 500 nm and assumed equivalent at 490 nm), and the fixed optical pathlength of the plate reader (l = 0.10 cm) using the Beer–Lambert law:

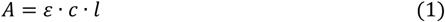

Initial velocities (*V*_0_) were obtained from the slope of the absorbance at 490 nm versus time during the initial linear phase of the assay and converted to concentration per unit time according to:

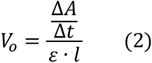

To determine the kinetic constants *K*_*m*_ and *V*_*max*_, assays were performed at multiple substrate concentrations and *V*_0_ values were fitted to the Michaelis–Menten equation:

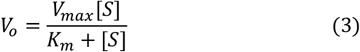

using the double reciprocal Lineweaver–Burk transformation, plotting 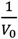 (where *V* is the initial velocity) versus 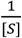 (where [*S*] is the nitrocefin concentration). *K*_*m*_ and *V*_*max*_ values are reported with standard error, and linear regressions were performed using GraphPad Prism 8 software.

For the estimation of the inhibition constant (*K*_*i*_) under the assumption of a competitive inhibition model, the following relationship was applied:

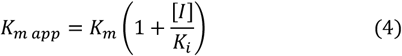

which rearranges to:

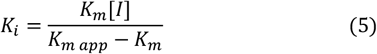

where *K*_*m app*_ corresponds to the apparent *K*_*m*_ value in the presence of the inhibitor, and [*I*] represents the inhibitor concentration, in this case, the antibody.

## Results

### Verification of the Expression of β-Lactamases and sd-Ab by Western Blot

The expression of the β-lactamases (BlaMab-2, KPC-2, VIM-2, OXA-48) was confirmed by SDS-PAGE and Western blot analysis. SDS-PAGE gels displayed bands corresponding to the expected molecular weights: BlaMab-2 at 33.7 kDa, KPC-2 at 33.2 kDa, VIM-2 at 31.4 kDa and OXA-48 at 33.4kDa, indicating proper expression in the *E. coli* BL21(DE3) system (Figure 2A). Western blot analysis using anti-His rabbit and anti-rabbit-ECL antibodies confirmed the presence of the target proteins with strong bands at the expected positions (Figure 2B).

**Figure 2.**
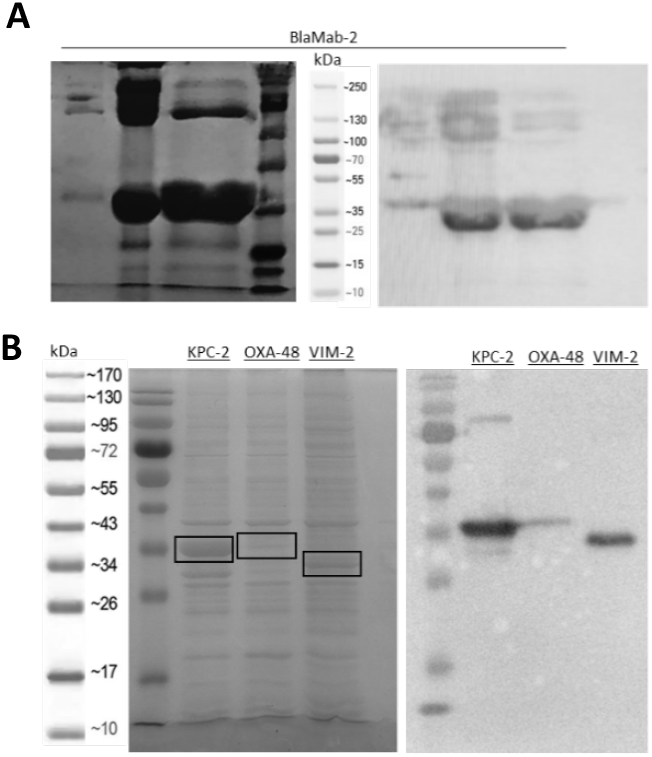
SDS-PAGE and Western blot analysis of β-lactamases. (A) SDS-PAGE electrophoresis gel of three clones of BlaMab-2 after affinity purification (left) and Western blot (anti-His) of the same clones (right). (B) SDS-PAGE electrophoresis of protein extracts from KPC-2, VIM-2 and OXA-48 expressing bacteria (left) and Western blot (anti-His) confirming the presence of the three β-lactamases (right).

### Expression and activity of β-Lactamases

The enzymatic activity of purified BlaMab-2 *in vitro* was assessed using nitrocefin hydrolysis assays. A significant increase in absorbance at 490 nm was observed over time, confirming that the enzyme was catalytically active. Higher nitrocefin concentrations resulted in increased hydrolysis rates, indicating substrate-dependent activity (Figure 3A). In addition, the enzymatic activity of the four β-lactamases (BlaMab-2, KPC-2, VIM-2, OXA-48) was evaluated in a cellular context using recombinant *E. coli* BL21(DE3) strains and a nitrocefin final concentration of 100 µM. Absorbance at 490nm increased over time for all of them at the same substrate concentration, confirming their functionality in whole cells (Figure 3B).

**Figure 3.**
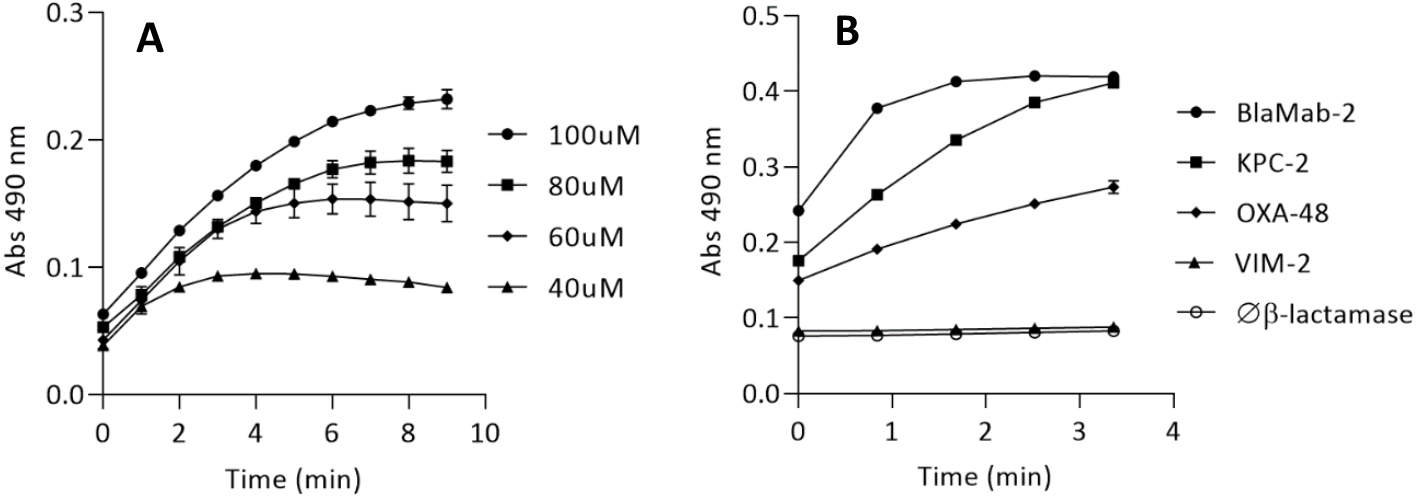
Enzymatic activity assays. (A) *In vitro* enzymatic activity of BlaMab-2 measured by the hydrolysis of nitrocefin at increasing concentrations at 490 nm. (B) *In vivo* activity of BlaMab-2, KPC-2 and OXA-48 in *E. coli* strains through detection of the absorbance at 490 nm produced by the hydrolysis of nitrocefin at a final concentration of 100 µM.

### Kinetic Parameters of β-Lactamases and Inhibition by B2 and B5

After obtaining the graphs and fitting the values to the Michaelis-Menten equation, K_m_, V_max_ and K_i_ values were obtained for purified BlaMab-2 in the presence and absence of sd-Abs B2, B5 (Table 1) and B7 (*Supplementary material*) *in vitro*. In the case of all three sd-Abs the K_m_ increased and V_max_ remained stable, suggesting competitive inhibition. B2 had a lower K_i_ than B5, but of the same order of magnitude. A notable observation was that, while B2 and B5 exhibited positive K_i_ values indicative of inhibition, B7 showed a negative K_i_ value (−3.7 ± 0.6).

**Table 1.** Enzyme activity constants K_m_, V_max_ and K_i_ of purified β-lactamase BlaMab-2 for the hydrolysis of nitrocefin *in vitro*, in the absence and presence of two sd-Ab inhibitors: B2 and B5.

| ∅ sd-Ab | | B2 | | | B5 | | |
| --- | --- | --- | --- | --- | --- | --- | --- |
| $K_m$ μM | $V_{max}$ μM/min | $K_{mi}$ μM | $V_{max}$ μM/min | $K_i$ μM | $K_{mi}$ μM | $V_{max}$ μM/min | $K_i$ μM |
| $45,3 \pm 2,5$ | $0,22 \pm 0,01$ | $64,4 \pm 2,2$ | $0,25 \pm 0,008$ | $2,4 \pm 0,08$ | $57,3 \pm 0,9$ | $0,23 \pm 0,03$ | $3,8 \pm 0,06$ |

Similarly, kinetic parameters were obtained for KPC-2, VIM-2, and OXA-48 in whole-cell assays. The addition of B2 and B5 increased K_m_ values, with B2 exhibiting stronger inhibition (Table 2).

**Table 2.** Enzyme activity constants K_m_, V_max_ and K_i_ of the β-lactamases BlaMab-2, KPC-2, VIM-2 and OXA-48 produced *in vivo* by *E. coli* strain BL21(DE3) for the hydrolysis of nitrocefin, in the absence and presence of the sd-Abs inhibitors B2 and B5.

| | ∅ sd-Ab | B2 | | B5 | |
| --- | --- | --- | --- | --- | --- |
| | $K_m$ μM | $K_{mi}$ μM | $K_i$ μM | $K_{mi}$ μM | $K_i$ μM |
| **BlaMab-2** | $48,029 \pm 1,28$ | $110,11 \pm 2,068$ | $1,52 \pm 0,028$ | $159,53 \pm 2,015$ | $5,90 \pm 0,074$ |
| **KPC-2** | $1,36 \pm 0,11$ | $120,91 \pm 2,73$ | $0,060 \pm 0,0014$ | $2,19 \pm 3,62$ | $0,12 \pm 0,0068$ |
| **VIM-2** | $4,35 \pm 0,035$ | $39,61 \pm 0,78$ | $2,90 \pm 0,057$ | $0,97 \pm 0,026$ | $68,92 \pm 1,85$ |
| **OXA-48** | $4,48 \pm 0,062$ | $6,62 \pm 0,16$ | $10,51 \pm 0,25$ | $2,26 \pm 0,058$ | $30,77 \pm 0,79$ |

## Discussion

### Heterologous Expression of β-lactamases and sd-Abs

The results from SDS-PAGE and Western blot analyses confirmed the successful expression and purification of β-lactamases in the recombinant *E. coli* BL21(DE3) system. Distinct protein bands corresponding to the expected molecular weights (BlaMab-2: 33.7 kDa; KPC-2: 33.2 kDa; VIM-2: 31.4 kDa; OXA-48: 33.4 kDa) were observed, validating appropriate expression of the constructs. The specificity of these bands was further verified through Western blot detection using anti-His antibodies. Together, these findings confirm that the recombinant proteins were correctly cloned and expressed in sufficient quantities, thereby providing a reliable source of material for subsequent *in vitro* and *in vivo* analyses.

### Expression and activity of β-Lactamases

The nitrocefin hydrolysis assays confirmed that the purified recombinant BlaMab-2 protein is catalytically active, as evidenced by the concentration-dependent increase in absorbance at 490 nm (Figure 3). Kinetic analysis yielded a K_m_ value of 45.3 ± 2.5, which is higher than those reported by Ramírez et al. in 2017 (29 ± 4) and Soroka et al. in 2014 (24 ± 7) for recombinant BlaMab expressed in *E. coli*, suggesting a lower substrate affinity in our preparation. While the kinetic parameters are broadly consistent with previous studies, the observed differences in K_m_ may reflect variations in expression systems, purification strategies, or assay conditions that influence protein folding and conformational stability, thereby affecting substrate interaction.

Nitrocefin hydrolysis assays performed in *E. coli* BL21(DE3) strains producing BlaMab-2, KPC-2, VIM-2, and OXA-48 also demonstrated increased absorbance at 490 nm (Figure 3), confirming that these β-lactamases are functionally expressed in the bacterial host. The kinetic parameters obtained are summarized in Table 3. For BlaMab-2, the in vivo K_m_ value (48.0 ± 1.3 μM) was consistent with that of the purified protein (45.3 ± 2.5 μM), suggesting that the cellular environment does not substantially affect substrate affinity. For KPC-2, VIM-2, and OXA-48, only in vitro K_m_ values for purified enzymes have been reported in the literature, preventing direct comparisons with the present in vivo data. Nevertheless, the lower K_m_ values obtained here may reflect differences in assay conditions and the impact of the cellular context, including expression system and protein folding within *E. coli* BL21(DE3).

**Table 3.** In vivo K_m_ values of recombinant β-lactamases and their inhibition by B2 and B5, compared with published in vitro data for nitrocefin hydrolysis.

| | $K_m$ μM (literature) | $K_m$ μM | $K_m$ with B2 μM | $K_m$ with B5 μM |
| --- | --- | --- | --- | --- |
| **BlaMab-2** | 24 ± 7.0<br>(Soroka et al., 2014) | 48,029 ± 1,28 | 110,11 ± 2,068 | 159,53 ± 2,015 |
| **KPC-2** | 45 ± 2.1<br>(Sun et al., 2024) | 1,36 ± 0,11 | 120,91 ± 2,73 | 2,19 ± 3,62 |
| **VIM-2** | 18<br>(Docquier et al., 2003) | 4,35 ± 0,035 | 39,61 ± 0,78 | 0,97 ± 0,026 |
| **OXA-48** | 23,9 ± 3.5<br>(Aertker et al., 2020) | 4,48 ± 0,062 | 6,62 ± 0,16 | 2,26 ± 0,058 |

### Evaluation of sd-Ab B2 and B5 as inhibitors

The study of new β-lactamase inhibitors is a promising and commonly adopted strategy to combat β-lactam antibiotic resistance (Zhou et al., 2022). In this study, we evaluated two sd-Abs, B2 and B5, against four clinically relevant β-lactamases. Both antibodies inhibited BlaMab-2, as reflected by increased K_m_ values while V_max_ remained unchanged, consistent with a competitive inhibition mechanism. Inhibition constants further highlighted their potency, with B2 consistently outperforming B5 across all tested enzymes, suggesting higher affinity for the active site and more efficient competition with the substrate.

In contrast, B7 produced a negative K_i_ (Supplementary Material), which is not meaningful in enzymatic kinetics and indicates it does not act as a genuine inhibitor under our conditions. Such results can arise from data-fitting limitations or from noncanonical interactions that enhance, rather than suppress, activity. One possibility is that B7 does not block substrate access directly but induces a conformational change that stabilizes BlaMab-2 or increases substrate affinity. This would manifest as an apparent activation rather than inhibition. Further structural and mechanistic studies are required to clarify whether B7 functions as an allosteric modulator rather than a competitive inhibitor.

In addition, the data summarised in Table 3 show an increase in K_m_ of the four proteins *in vivo* in the presence of B2, indicating a significant decrease in substrate affinity due to its inhibitory effect. This decrease in affinity suggests a competitive inhibition mechanism, where the inhibitor binds directly to the active site of the enzyme, blocking substrate binding. Consistently, the in vivo inhibition assays demonstrated that B2 significantly reduced the enzymatic activity of BlaMab-2, KPC-2, VIM-2, and OXA-48, reinforcing its broad-spectrum inhibitory potential. However, the case of VIM-2 and OXA-48 in the presence of sd-Ab B5 is striking. Unlike B2, B5 led to a decrease in K_m_ for these enzymes, suggesting an increase in substrate affinity. This pattern is characteristic of an uncompetitive inhibition mechanism, where the inhibitor preferentially binds to the enzyme-substrate complex rather than the free enzyme, stabilizing an inactive state. Such a mode of inhibition is less common but can offer distinct therapeutic advantages, particularly in conditions where substrate concentrations are high, reducing the effectiveness of competitive inhibitors.

The inhibitory activity of the sd-Abs B2 and B5 was assessed by comparing their K_i_ values with those reported for established β-lactamase inhibitors (Table 4). Conventional inhibitors such as clavulanic acid, tazobactam, sulbactam, and avibactam are increasingly compromised by resistant β-lactamases, underscoring the need for new strategies. In this study, both sd-Abs demonstrated highly favorable K_i_ values relative to those previously described. In the case of BlaMab-2, inhibition by sd-Ab B2 yielded a K_i_ of the same order of magnitude as clavulanic acid (Ramírez et al., 2017), while sd-Ab B5 achieved a notably lower value than relebactam (Dousa et al., 2020).

**Table 4.** Comparison of K_i_ values of sd-Ab B2 and B5 for the inhibition of BlaMab-2, KPC-2, VIM-2 and OXA-48 *in vivo* with those reported in the literature.

| | $K_i$ B2 $\mu\text{M}$ | $K_i$ B5 $\mu\text{M}$ | $K_i \mu\text{M}$ (literature) | | Ref. |
| --- | --- | --- | --- | --- | --- |
| **BlaMab-2** | $1,52 \pm 0,028$ | $5,90 \pm 0,074$ | Clavulanic acid | $2.3 \pm 0.7$ | (Ramírez et al., 2017) |
| | | | Durlobactam | $(4.0 \pm 0.8) \times 10^{-3}$ | (Dousa et al., 2022) |
| | | | Avibactam | $0.30 \pm 0.03$ | (Dousa et al., 2020) |
| | | | Relebactam | $136 \pm 14$ | (Dousa et al., 2020) |
| **KPC-2** | $0,060 \pm 0,0014$ | $0,12 \pm 0,0068$ | Clavulanic acid | 29.96 | (Khan et al., 2014) |
|  |  |  | Sulbactam | 23.21 | (Khan et al., 2014) |
|  |  |  | Tazobactam | 21.61 | (Khan et al., 2014) |
| | | | Taniborbactam | $0.004 \pm 0.001$ | (Hamrick et al., 2020) |
| | | | Avibactam | $0.0056 \pm 0.0007$ | (Hamrick et al., 2020) |
| | | | Vaborbactam | $0.022 \pm 0.002$ | (Hamrick et al., 2020) |
| **VIM-2** | $2,90 \pm 0,057$ | $68,92 \pm 1,85$ | Mitoxantrone | $1,5 \pm 0,2$ | (Minond et al., 2009) |
| | | | Taniborbactam | $0.019 \pm 0.001$ | (Hamrick et al., 2020) |
| **OXA-48** | $10,51 \pm 0,25$ | $30,77 \pm 0,79$ | Taniborbactam | $0.35 \pm 0.007$ | (Hamrick et al., 2020) |
| | | | Avibactam | $0.26 \pm 0.005$ | (Hamrick et al., 2020) |
| | | | Vaborbactam | $0.35 \pm 0.007$ | (Hamrick et al., 2020) |

For KPC-2, sd-Ab B2 displayed a K_i_ value comparable to that of vaborbactam (Hamrick et al., 2020). Moreover, the K_i_ values for B2 and B5, were three and two orders of magnitude lower, respectively, than those reported by Khan et al. in 2014 for clavulanic acid, sulbactam and tazobactam. In the case of VIM-2, the K_i_ obtained in this study with B2 was slightly higher than that reported by Minond et al. in 2009 with mitoxantrone, yet remained within a range supporting its potential as a VIM-2 inhibitor. For OXA-48, B2 showed K_i_ values two orders of magnitude higher than those reported in the literature for taniborbactam, avibactam and varbobactam (Hamrick et al., 2020). As with VIM-2, these results suggest that B2 may still serve as a valuable inhibitor, particularly under conditions where resistance compromises existing compounds. Overall, both sd-Abs demonstrated strong inhibitory activity, B2 consistently outperformed B5 as the most promising sd-Ab inhibitor due to its strong and consistent competitive inhibition profile across multiple β-lactamases. In addition, it also shows better inhibition values (K_i_) than B5 for all four proteins. B5, while less potent, remains a promising candidate warranting further evaluation.

Despite the promising inhibitory activity observed in this study, several challenges must be addressed before sd-Abs can be developed as therapeutic agents. First, further structural and biophysical studies, such as X-ray crystallography or cryo-EM, are necessary to elucidate the precise binding interactions between sd-Abs and β-lactamases. This information would facilitate the rational design of improved inhibitors with enhanced affinity and specificity. Second, *in vivo* studies are necessary to assess the pharmacokinetics, stability, and therapeutic efficacy of sd-Abs in infection models. Finally, large-scale production strategies must be optimized to enable cost-effective manufacturing for clinical use. Overall, the differential behaviour of B2, B5, and B7 underscores the complexity of β-lactamase inhibition by sd-Abs and highlights the necessity of further characterization to fully exploit their therapeutic potential.

## Conclusions

This study confirmed the enzymatic activity of four recombinant β-lactamases produced in *E. coli* BL21(DE3): BlaMab-2 from *Mycobacterium abscessus*, KPC-2 and OXA-48 from *Klebsiella pneumoniae* and VIM-2 from *Pseudomonas aeruginosa*. The sd-Abs B2 and B5, generated against BlaMab-2, effectively inhibited this enzyme both in vitro and in vivo via competitive inhibition. Notably, B2 also inhibited KPC-2, VIM-2, and OXA-48 competitively, while B5 displayed an acompetitive pattern for VIM-2 and OXA-48.

Overall, our results highlight the importance of exploring new therapeutic strategies based on sd-Abs to combat the growing threat of antibiotic resistance. These findings open up the possibility of using a broader range of β-lactam antibiotics, in combination with sd-Ab B2 and B5, expanding their spectrum of action and enhancing therapeutic efficacy against multidrug-resistant bacterial infections.

## Acknowledgments

This work was funded by a grant from Ministerio de Ciencia e Innovación PID2022-136371OB-I00 to AP-A and MP.

## Supplementary material

**Table 5.** Enzyme activity constants K_m_, V_max_ and K_i_ of purified β-lactamase BlaMab-2 for the hydrolysis of nitrocefin *in vitro*, in the absence and presence of the sd-Ab B7.

| **Ø sd-Ab** |  | **B7** |  |  |
| --- | --- | --- | --- | --- |
| $K_m \mu M$ | $V_{max} \mu M/min$ | $K_{mi} \mu M$ | $V_{max} \mu M/min$ | $K_i \mu M$ |
| $45,3 \pm 2,5$ | $0,22 \pm 0,01$ | $57,3 \pm 0,9$ | $0,18 \pm 0,03$ | $-3,7 \pm 0,6$ |

